# Network analysis reveals the molecular bases of statin pleiotropy that vary with genetic background

**DOI:** 10.1101/2022.07.17.500365

**Authors:** Cintya E. del Rio Hernandez, Lani J. Campbell, Paul H. Atkinson, Andrew B. Munkacsi

## Abstract

Many approved drugs are pleiotropic, for example statins, whose main cholesterol lowering activity is complemented by anticancer and pro-diabetogenic mechanisms involving poorly characterized genetic interaction networks. We investigated these using the *Saccharomyces cerevisiae* genetic model where most genetic interactions known are limited to the statin-sensitive S288C genetic background. We therefore broadened our approach by investigating gene interactions to include two statin-resistant UWOPS87-2421 and Y55 genetic backgrounds. Networks were functionally focused by selection of *HMG1* and *BTS1* mevalonate pathway genes for detecting genetic interactions. Networks, multi-layered by genetic background, were analysed for modifying key genes using network centrality (degree, betweenness, closeness), pathway enrichment, functional community modules and gene ontology. Statin treatment induces the unfolded protein response and we found modifying genes related to dysregulated endocytosis and autophagic cell death. To translate results to human cells, human orthologues were searched for other drugs targets, thus identifying candidates for synergistic anticancer bioactivity.

## Introduction

Since their discovery more than 40 years ago, statins (Endo et al. 1976a) have saved millions of lives via cholesterol reduction and prevention of cardiovascular disease by competitive inhibition of the rate-limiting 3-hydroxy-3-methyl-glutaryl-coenzyme A reductase (HMGCR) enzyme in the mevalonate pathway (Fig 1) (Endo et al. 1976b, Goldstein and Brown 1973). But statins, like many drugs, are pleiotropic and affect other pathways including those related to diabetes and tumorigenesis (Betteridge and Carmena 2016, Juarez and Fruman 2021, King et al. 2022, Göbel et al. 2020; Hart et al. 2015). Such pleiotropy may be exploited to investigate other useful properties of such drugs. Pleiotropic properties of drugs are often the consequence of complex gene network dynamics downstream of the primary target.

**Figure 1.**
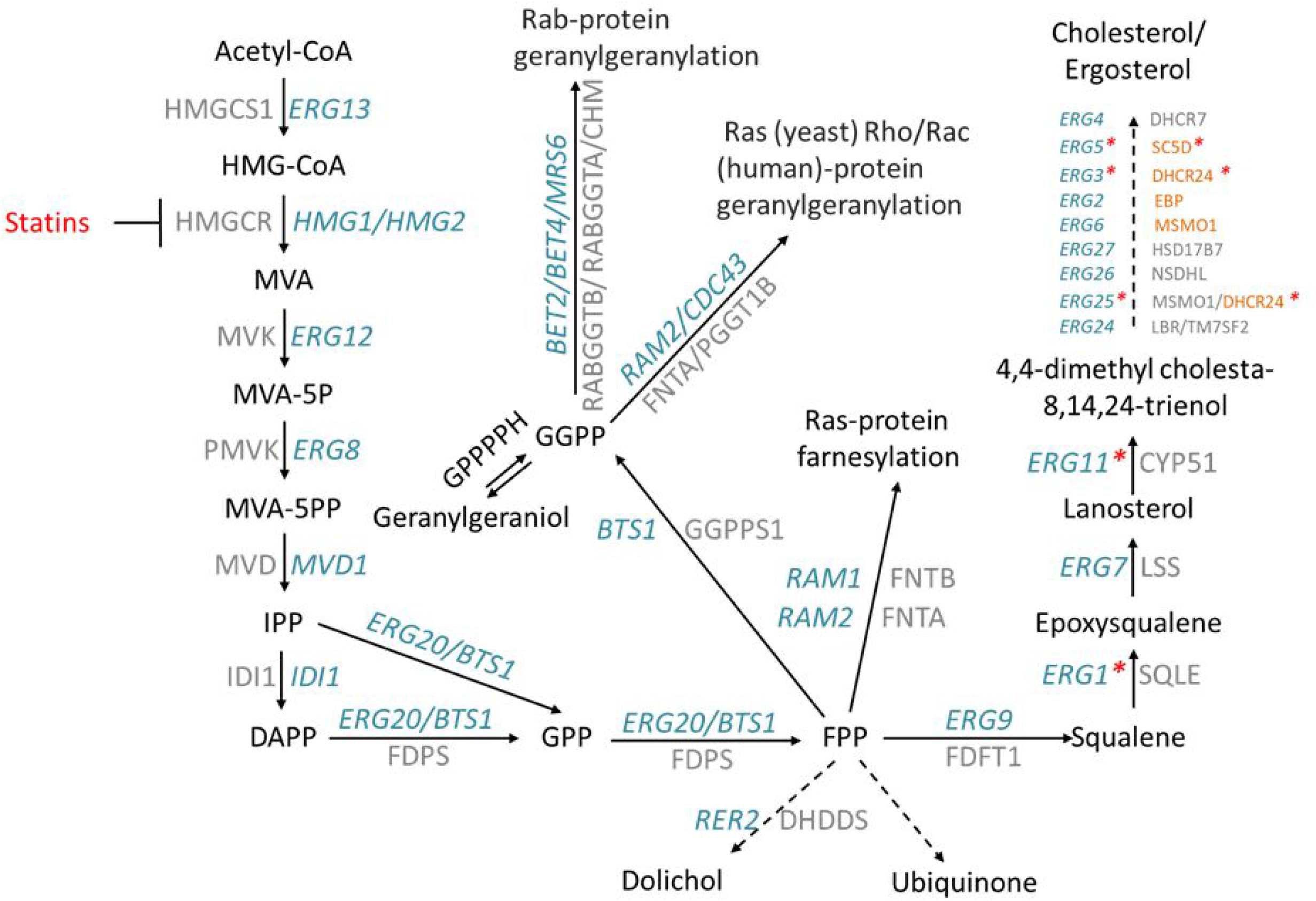
Statins inhibit the synthesis of HMGCR and downstream products in the mevalonate pathway. Statins are competitive inhibitors of HMGCR encoded by *HMG1* and *HMG2* in yeast and HMGCR in humans. A critical step in the mevalonate pathway is mediated by the enzyme geranylgeranyl diphosphate synthase (encoded by *BTS1* in yeast and GGPPS1 in humans), where the main ergosterol/cholesterol-synthesis pathway branches off to synthesise other fundamental cellular components for isoprenylation of small GTPases. Genes in blue are yeast genes and genes in grey are their human orthologues. Red asterisks in yeast genes indicate oxygen-dependent steps of the pathway. Human genes in orange at the end of the cholesterol pathway are less conserved with yeast and do not correspond to the yeast gene to the left.

Yeast *Saccharomyces cerevisiae* (*S. cerevisiae*) is an established eukaryote model organism (Botstein et al. 1997) where about 70% of its genes show high conservation with humans not only in sequence but also in biological function (Goffeau et al. 1996). This was cleanly illustrated with humanisation of yeast (*i.e*., expression of human genes in yeast) where only 20% amino acid identity was required for human genes to complement the deletion of orthologous yeast genes (Kachroo et al. 2015). Relevant to this study, human HMGCR restored the viability of yeast lacking its two paralogue genes, *HMG1* and *HMG2* (Basson et al. 1988). Indeed, many steps of the mevalonate pathway were originally elucidated in yeast (Bloch 1965; Hampton and Rine 1994). Because of its genetic tractability, it is a powerful aid for the study of the mevalonate pathway (Bloch 1965; Hampton and Rine 1994), cancer cell biology (Ahangari et al. 2020; Guan et al. 2019; Mullen et al. 2016) and complex phenotypes in general (Busby et al. 2019; Ferreira et al. 2019; Hartwell et al. 1997; Munkacsi et al. 2011; Simon 2001).

Complexity may be investigated by genetic interactions involving epistasis (Phillips 1998), which measures functionality shared by the interacting pairs of genes. In yeast, interactions may be scored in high-throughput screens called synthetic genetic array (SGA) analysis that measure colony size phenotype changes exerted in pairs of double gene deletion strains (Tong et al. 2001; Boone et al; 2007), or by a gene deletion paired with a gene-product inhibitory drug (Parsons et al, 2006). The tractability of yeast genetics allowed genome-wide cataloguing of genetic interactions that are called synthetic lethal when a double mutant exhibits no growth or synthetic sick when the double mutant exhibits reduced growth (Dobzhansky 1946, Costanzo et al. 2019).

From these synthetic lethal and synthetic sick interactions, gene networks have been assembled representing 5.4 million interactions in the S288C genetic background (Costanzo et al. 2010; Costanzo et al. 2016). Networks in turn may be analysed for key genes using graph centrality metrics (Boccaletti et al. 2014, Dong and Horvath 2007, Newman 2005, Brandes 2001), and here we applied such methodology to statin pleiotropy. We had at hand three libraries of yeast genome-wide deletion strains constructed in three different genetic backgrounds S288C, Y55, and UWOPS87-2421 (hereafter referred as UWOPS87) (Busby et al, 2019; Galardini et al. 2019), allowing us to additionally characterise statin pleiotropy by genetic background.

## Results

The overall scheme of our paper (Fig 2) is to elucidate the mevalonate pathway-specific genetic interactions integral to statin bioactivity. Using SGA methodology (Tong et al. 2001), we generated 25,800 double deletion yeast strains, each lacking a gene in the statin pathway and a second gene in the yeast genome of statin-susceptible and statin-resistant genetic backgrounds since cholesterol-lowering activities of statins vary among individuals (Ahangari et al. 2020; Guan et al. 2019). The genes within the mevalonate pathway investigated were *HMG1*, the predominantly active target of atorvastatin under aerobic conditions (about 80% of the activity compared to its paralogue *HMG2* (Basson et al. 1986)), and *BTS1*, the mediator of the off-branch pathway from the main ergosterol synthesis pathway to isoprenylation of GTPases. The double deletion mutants were treated with atorvastatin and hypersensitive mutants were compiled into multi-layered networks. Topology centrality metrics and functional enrichment in chemical genetic interaction networks were used to identify key genes and cellular processes regulating statin activity, which by definition are candidate targets to use in combination with statins to enhance its anticancer activity.

**Figure 2:**
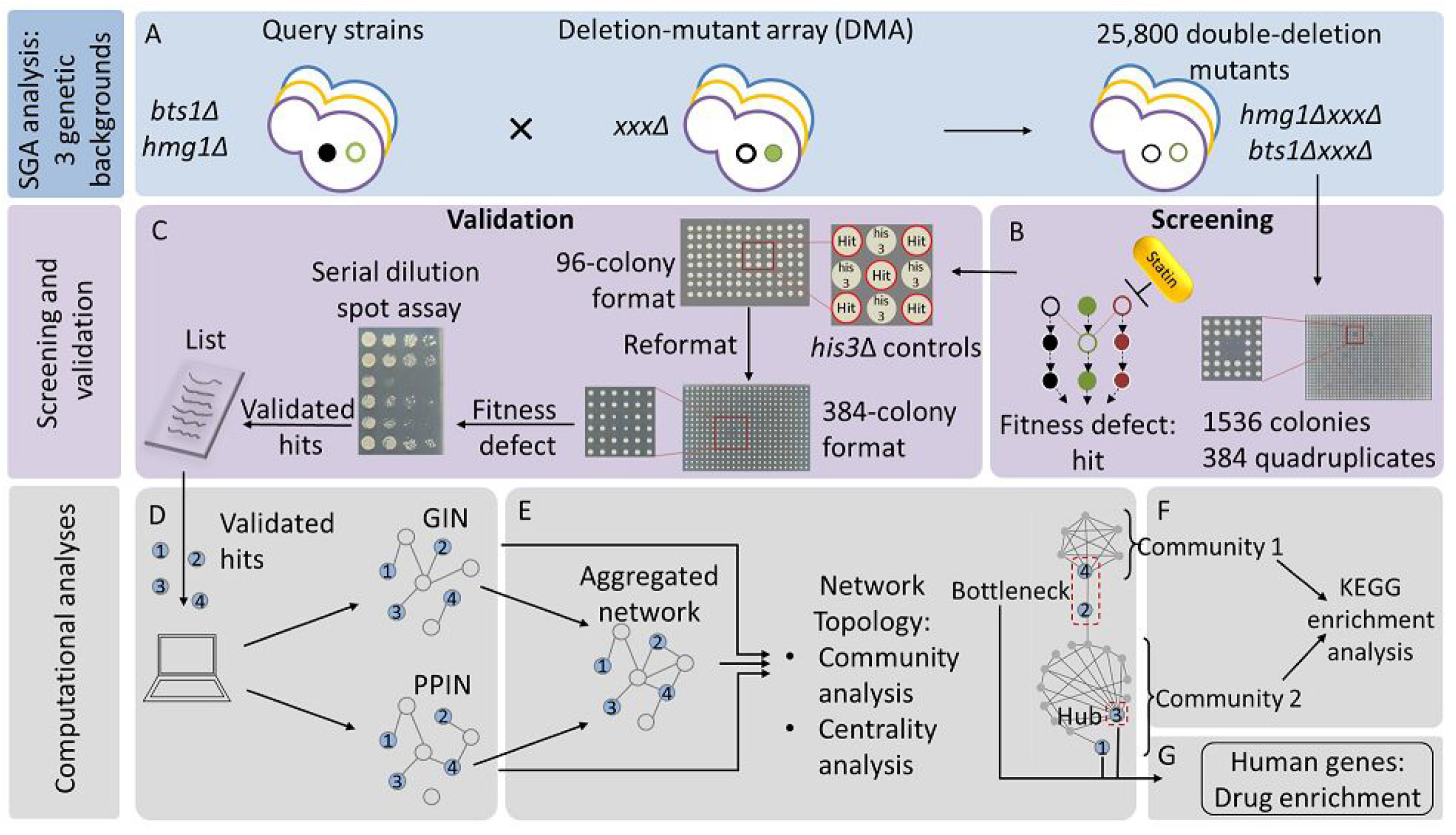
Flow diagram for the methods used to identify interactions and pathways behind atorvastatin pleiotropy in three genetic backgrounds. Single deletion mutant query strains were constructed (A) (deletion mutant genes depicted as empty circles) as models to investigate the anticancer activity of atorvastatin (*hmg1*Δ and *bts1*Δ) in three yeast genetic backgrounds (S288C, UWOPS87 and Y55 indicated here as purple, yellow and blue), and mated against DMAs of the same genetic backgrounds to generate 25,800 double deletion mutants in 1536-colony format (384 quadruplicate colonies per agar plate). These mutants were treated with atorvastatin (B) and screened to identify fitness defects that would reveal epistatic interactions as measured by decreased colony size. Atorvastatin-hypersensitive double mutants were then validated in two steps (C). First, hypersensitive mutants were formatted in 96-colony format plates with each one surrounded by *his3*Δ strains for growth control. These plates were then reformatted to 384-colony format (96 quadruplicate colonies) and screened again with atorvastatin. Colonies that showed fitness defects were selected for the second step, which consisted of serial dilution spot assays. Double deletion mutants that showed growth inhibition in the latter were considered as validated interactions and used as input to create genetic (GIN) and protein-protein (PPIN) interaction networks (D). GINs and PPINs were multi-layered in one network (E) per genetic background and subjected to network topology analyses. The network centrality metrics pinpointed bottleneck and hub genes of high biological relevance. The communities of genes identified through network modularity (F) were analysed through a KEGG enrichment analysis to distinguish key metabolic pathways. Human orthologues of the key yeast genes were used in a search for drug enrichment (G) to identify potential combination therapies to enhance the anticancer activity of atorvastatin.

### Screening for statin-specific epistasis in genome-wide deletion libraries in three genetic backgrounds

In order to measure the chemical genetic effects of atorvastatin and the combined *hmg1Δ xxxΔ* and *bts1Δ xxxΔ* double gene deletions, it was necessary to ensure atorvastatin was not present in excess so statin-effect and double mutant-effect could be distinguished. To achieve this, we determined the concentration of atorvastatin that reduced the growth of single deletion mutant *hmg1Δ* and *bts1Δ* to 70% of normal growth in the S288C, Y55, UWOPS87 genetic background strains. Accordingly, we separately deleted the *HMG1* and *BTS1* genes through PCR-directed mutagenesis and homologous recombination in the three backgrounds and then treated with atorvastatin to characterise the toxicity range of concentrations of the drug (Fig 3A). All three genetic backgrounds showed the same sensitivity when *HMG1* was deleted (*i.e*., synthetic sick at 5 μM atorvastatin, synthetic lethal at 50 μM atorvastatin), probably because all the backgrounds are equally reliant on *HMG1* to cope with atorvastatin. Contrastingly, when *BTS1* was deleted in S288C, synthetic lethality occurred in 1 μM atorvastatin, while the same concentration exerted only a mild fitness defect in UWOPS87 and Y55. This may be because the downstream *BTS1* mediates several branches from the mevalonate pathway, possibly providing background-specific statin resistance pathways.

**Figure 3.**
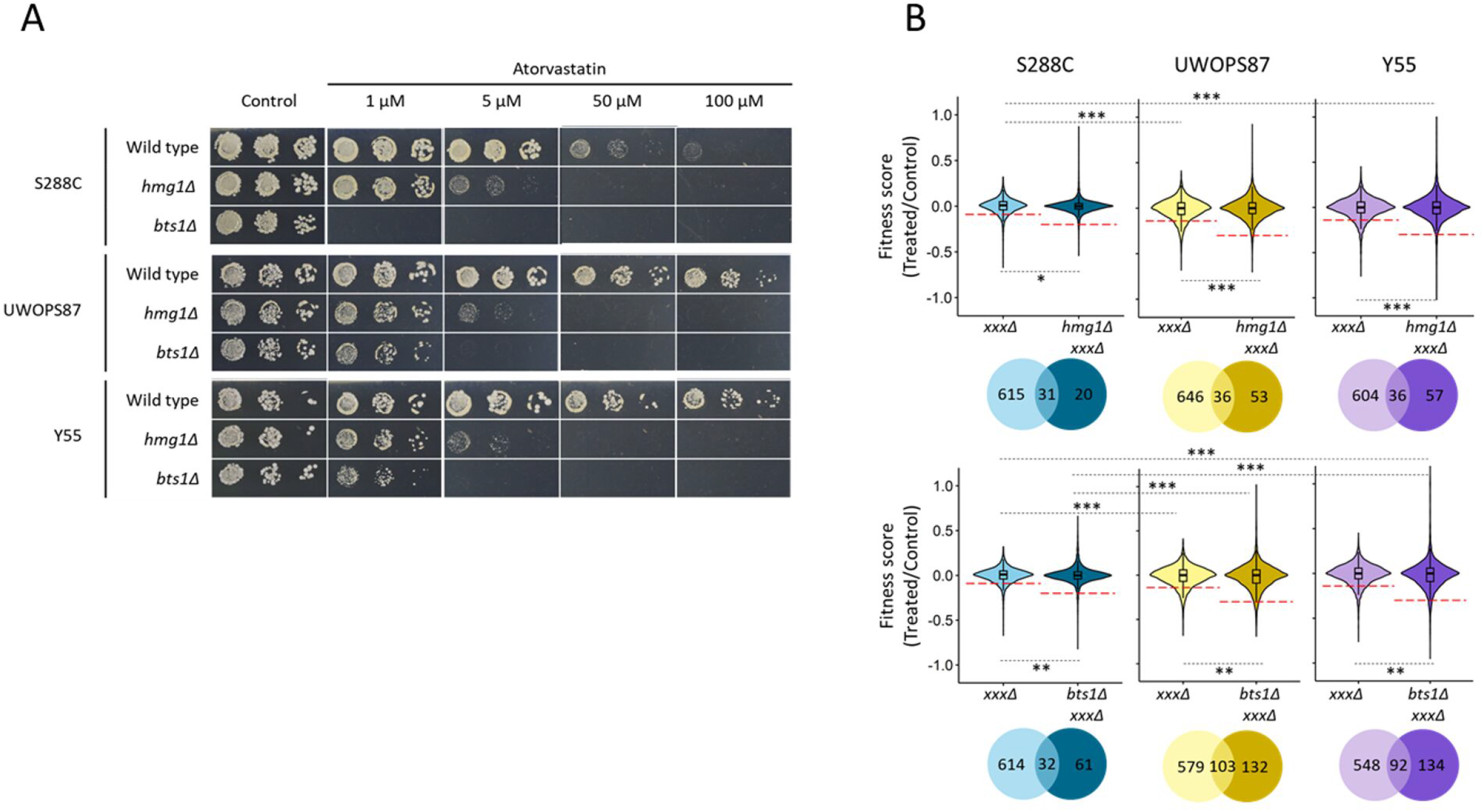
Atorvastatin sensitivity confers similar synthetic sickness/lethality in *HMG1*-deleted strains and varies in *BTS1-*deleted strains in three genetic backgrounds. (A) Haploid cells deficient of *HMG1* or *BTS1* and their wild types in three genetic backgrounds were pinned on increasing concentrations of atorvastatin in serial dilution and incubated for 2 days at 30°C. (B) Violin plot distributions of average fitness of 12,900 strains as measured by colony sizes (n = 4) of *xxxΔ* and *hmg1Δ xxxΔ* (upper panel) as well as *xxxΔ* and *bts1Δ xxxΔ* (lower panel) where positive scores represent increased fitness and negative scores represent decreased fitness. The red dashed lines indicate the score cut-off values selected for validation in independent assays for double deletions that did not overlap with the *xxxΔ* single deletions. Venn diagrams visualise the overlap in the number of genes below the cut-off lines. Statistical differences were evaluated with a Student’s *t*-test (*, *P* < 0.05; **, *P* < 0.01; ***, *P* < 0.001).

We then investigated atorvastatin-specific triple mutant chemical genetic epistasis in our three different genetic background (S288C, Y55, and UWOPS87) double deletion libraries using *hmg1Δ and bts1Δ* as SGA query strains. Thus the 70% (IC_30_) concentration in all libraries: *hmg1Δ xxxΔ, bts1Δ xxxΔ* and *xxxΔ* was chosen from the range 0.2 to 64 μM for *hmg1Δ xxxΔ*, 0.01 to 64 μM for *bts1Δ xxxΔ* and 10 to 320 μM for *xxxΔ*. Using this information, *hmg1Δ xxxΔ* double deletions were thus screened at 0.8 μM atorvastatin, *bts1Δ xxxΔ* double deletions in S288C were screened at 0.05 μM and *bts1Δ xxxΔ* double deletions in Y55 and UWOPS87 at 0.5uM. The single deletion library *xxxΔ* control was screened with 9 μM for S288C, 10 μM for UWOPS87 and 35 μM for Y55. Violin plots showed that the distribution of scored colony sizes did not differ among the three genetic backgrounds when *HMG1* was deleted, but it did differ between S288C and the resistant genetic backgrounds when *bts1* was deleted (Fig 3B), thus adding evidence to our observations above that all three backgrounds are equally reliant on *HMG1*, but downstream *BTS1*-mediated pathway branches provide background specific resistance to atorvastatin.

### Experimental validation of atorvastatin-hypersensitive double deletion mutants

High-throughput screening experiments in high-density formats tend to suffer from false positive and false negative noisy data. To validate the atorvastatin-hypersensitive interactions, first we established a cut-off for the scored colonies (pixel-based colony size scored values assigned in SGAtools via Gitter (Wagih and Parts 2014)) of 3 standard deviations below the median for *hmg1Δ*strains and of 2.5 standard deviations below the median for *bts1Δ* strains (thus scores below −0.2 for S288C and below −0.3 for UWOPS87 and Y55 were considered genuine hypersensitive mutants). Strains that were sensitive in single and double deletion mutants in the presence of statins were excluded from further analysis (Fig 3B). Using these cut-off criteria, we found atorvastatin-specific interactions in 20, 53 and 57 *hmg1Δ xxxΔ* strains for S288C, UWOPS87 and Y55, respectively. Likewise for *bts1Δ xxxΔ* strains, there were 61, 132 and 134 atorvastatin-specific interactions in S288C, UWOPS87 and Y55, respectively. Atorvastatin-specific interactions in *hmg1Δ xxxΔ* (6 interactions in S288C, 8 interactions in Y55 and 11 interactions for UWOP287) and *bts1Δ xxxΔ* (7 interactions in S288C, 12 interactions in Y55 and 15 interactions in UWOP287) were then individually validated in a second step by plating in 384-colony quadruplicate format to confirm atorvastatin-sensitivity, followed by confirmation in serial spot dilution assays (Figs 4 and 5). Of the 40 yeast genes identified, 29 of them have human orthologues that have been previously annotated to cancer, 21 to diabetes, 10 to myopathies, 2 to rhabdomyolysis and 8 are known targets of statins (Table EV1).

**Figure 4.**
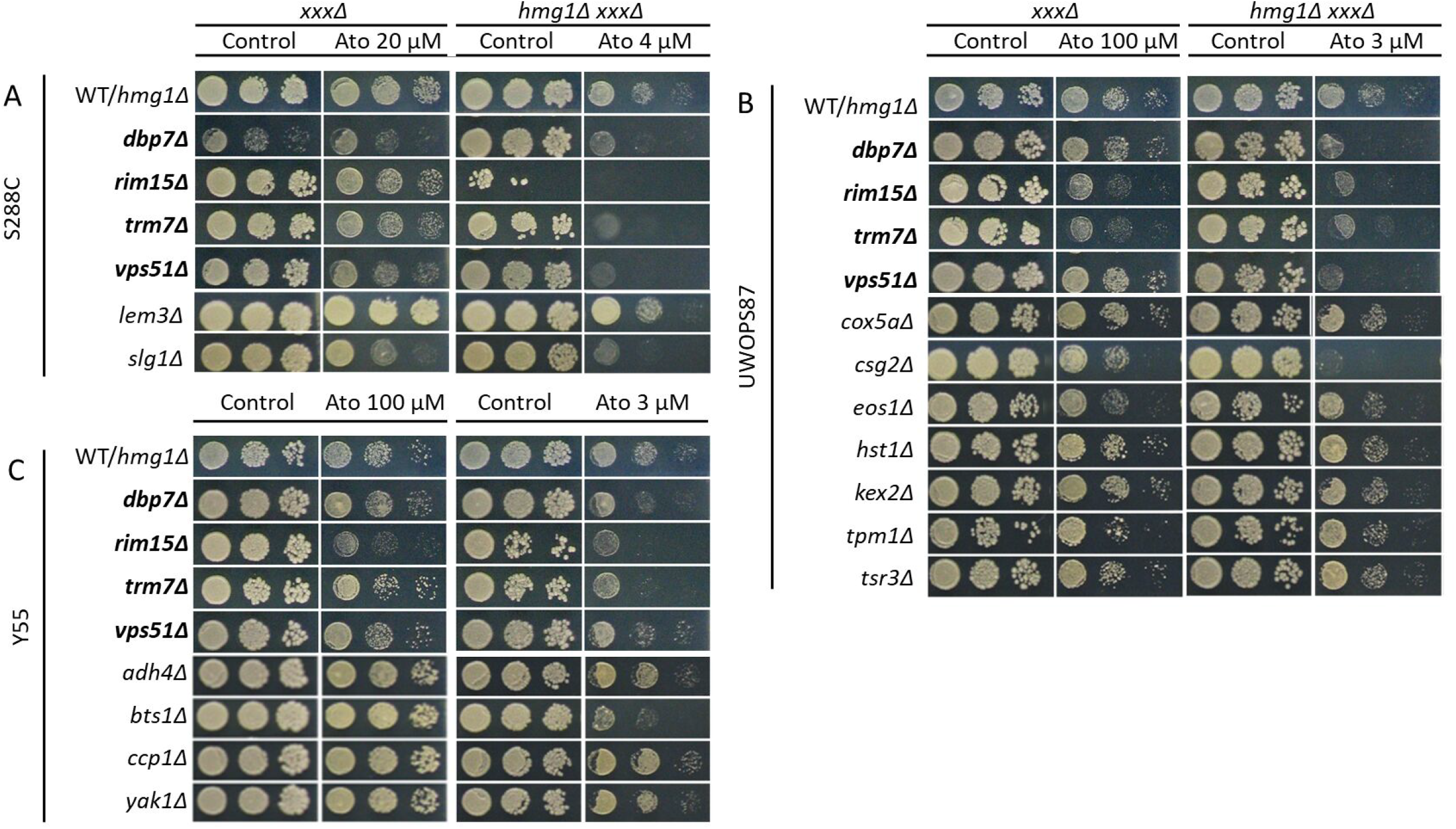
Four *hmg1Δ xxxΔ* double deletions were hypersensitive to atorvastatin treatment in three genetic backgrounds while others depend on genetic background. Haploid cells derived from SGA analyses and DMA libraries were pinned on SC with or without supplementation of atorvastatin in serial dilution and incubated for 2 days at 30°C. Shown here are deletions of genes that enhanced sensitivity to atorvastatin treatment in (A) S288C; (B) UWOPS87; and (C) Y55 genetic backgrounds. *WT/hmg1Δ* panel refers to either the non-mutated wild types (WT) for the *xxxΔ* strain panels or the *hmg1Δ* single deletions for the *hmg1Δ xxxΔ* double deletion strain panels. Gene deletions in bold indicate interactions overlapping in three genetic backgrounds.

**Figure 5.**
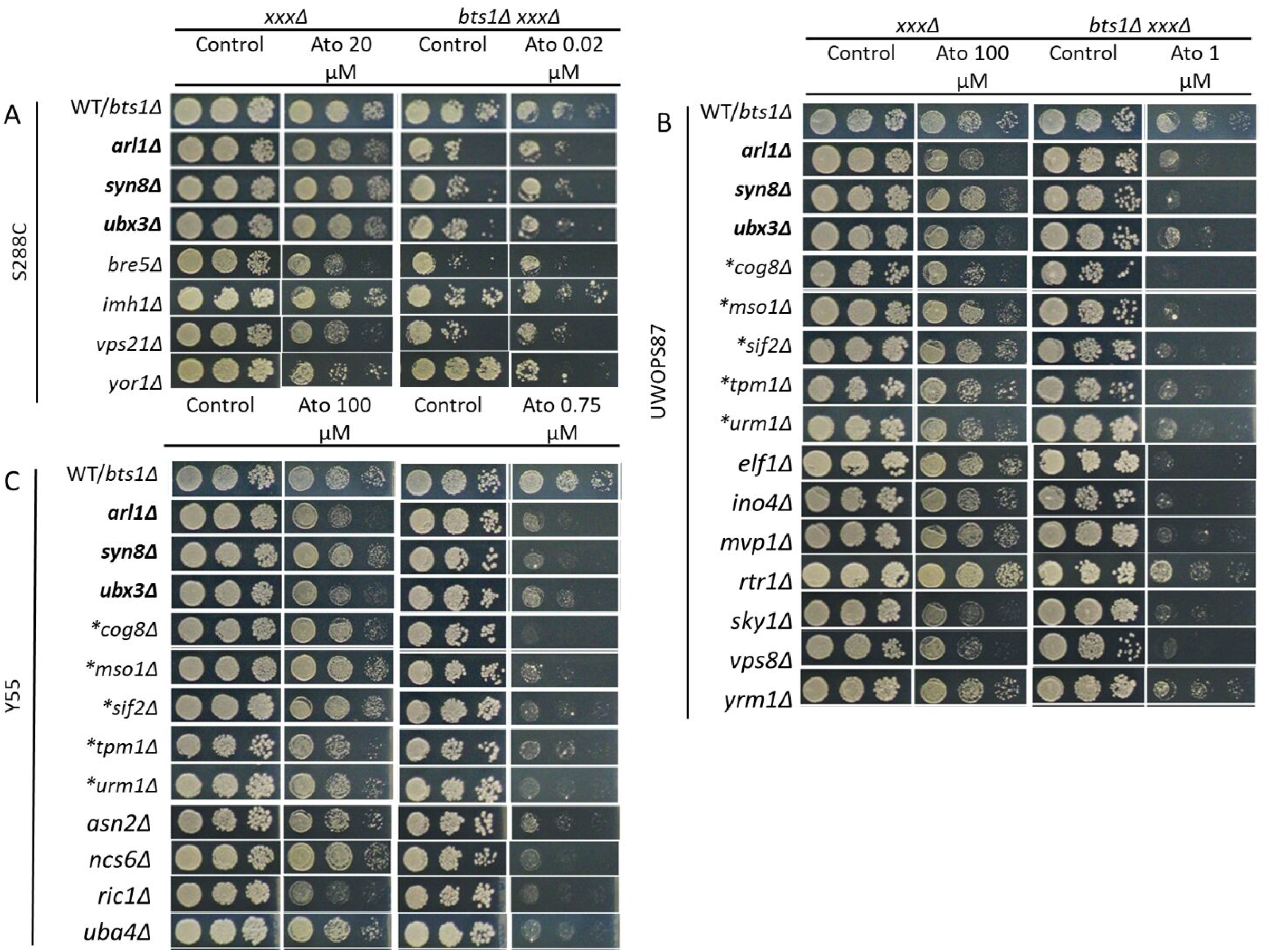
Eight *bts1Δ xxxΔ* double deletions were hypersensitive to atorvastatin treatment in at least two genetic backgrounds while others depend on the genetic background. Haploid cells derived from SGA analyses and DMA libraries were pinned on SC with or without supplementation of atorvastatin in serial dilution and incubated for 2 days at 30°C. Shown here are deletions of genes that enhanced sensitivity to atorvastatin treatment in (A) S288C; (B) UWOPS87; and (C) Y55 genetic backgrounds. *WT/bts1Δ* refers to either the non-mutated wild type (WT) for the *xxxΔ* strain panels or the *bts1Δ* single deletion for the *bts1Δ xxxΔ* double deletion strain panels. Gene deletions in bold indicate interactions overlapping in three genetic backgrounds, asterisks indicate interactions overlapping in two genetic backgrounds.

### Construction of multi-layered gene-gene and protein-protein interaction networks

Network centrality analyses are often performed in single-layer networks, that is, connections between nodes based on one type of functional relationship. Assessing multi-layer networks has now expanded the usefulness of centrality analyses by analysing two or more layers of interactions of different types of data (Wang et al. 2017). Similar to a single-layer network, albeit just more complex, multi-layered networks are basically n-dimensional matrices or tensors that can be investigated using graph mathematical methodologies.

We assembled multi-layered networks from the 17 and 23 genes (Figs 6 and 7) that were validated to be interactive with *HMG1* and *BTS1*, respectively, where the list of validated genes was augmented (pathlength 2) with genetic interaction networks (GINs) (Warde-Farley et al. 2010) and protein-protein interaction networks (PPINs) (Szklarczyk et al. 2015, Xia et al. 2015; Zhou et al. 2019). Thus, we identified and visualised GINs and PPINs specific to each type of interaction and each genetic background (Figs 6–7, EV1-EV2). The numbers of nodes and edges for GINs or PPINs were generally lower than those of the multi-layered network (Table 1; Table EV2), demonstrating that the connectivity of multi-layered networks was more robust than that of GINs and PPINs alone.

**Figure 6.**
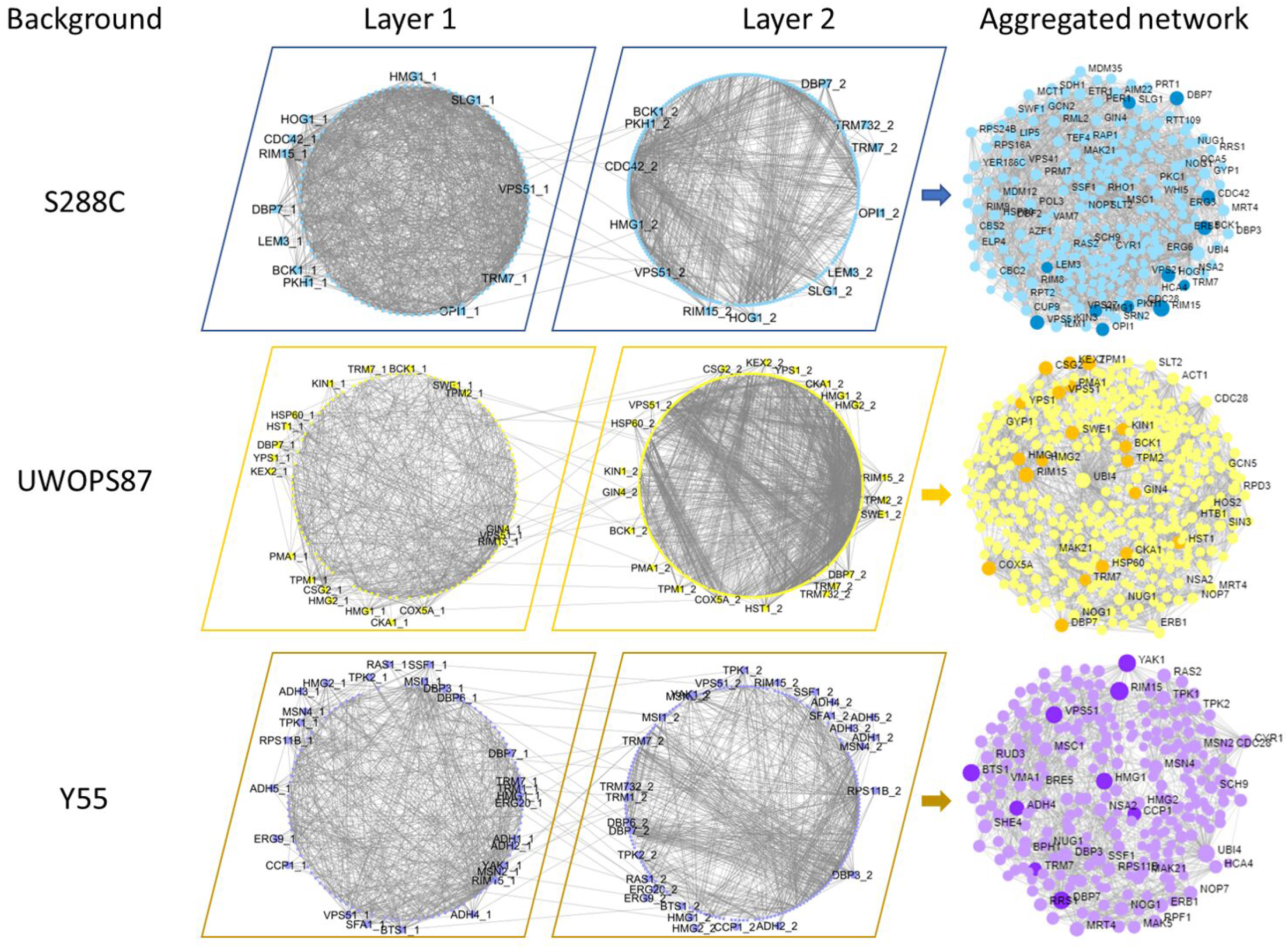
Multi-layer networks derived from atorvastatin-sensitive *hmg1Δ xxxΔ* interactions. GINs (Layer 1), PPINs (Layer 2) and the edges between them were integrated in an multi-layered network using TimeNexus. Edges between layers connect overlapping nodes in the two layers and the genes linking these edges are shown in the periphery of circular networks. Darker nodes in multi-layered networks are validated atorvastatin-sensitive interactions.

**Figure 7.**
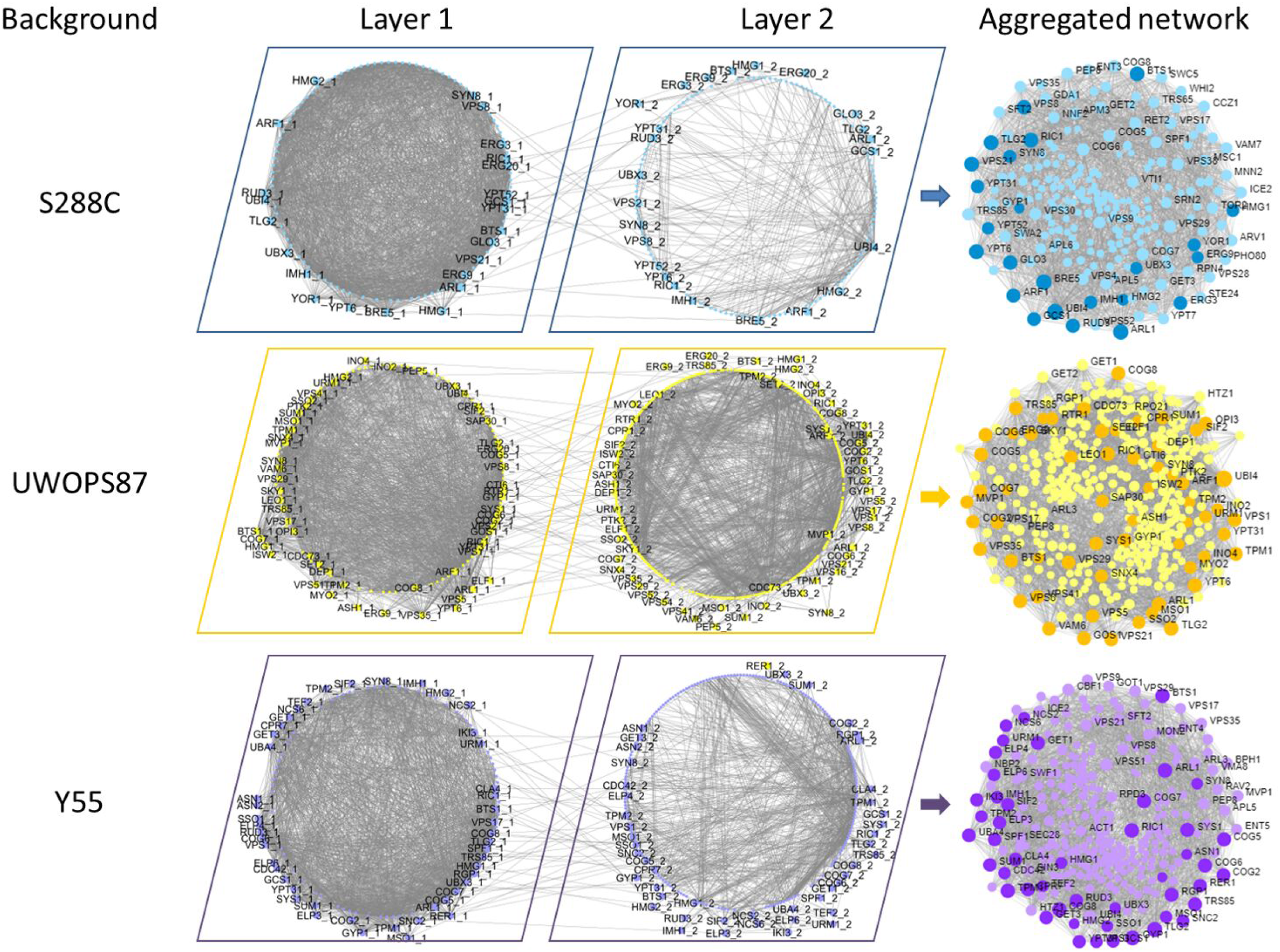
Multi-layer networks derived from atorvastatin-sensitive *bts1Δ xxxΔ* interactions. GINs (Layer 1), PPINs (Layer 2) and the edges between them were integrated in an multi-layered network using TimeNexus. Edges between layers connect overlapping nodes in the two layers and the genes linking these edges are shown in the periphery of circular networks. Darker nodes in multi-layered networks are validated atorvastatin-sensitive interactions.

**Table 1.** Top betweenness centrality measurements for multi-layer networks in three genetic backgrounds

| Query gene | Genetic Background | Nodes/ edges | Gene Name | Betweenness |  | Closeness |  | Degree |  |
| --- | --- | --- | --- | --- | --- | --- | --- | --- | --- |
|  |  |  |  | Score | Rank | Score | Rank | Score | Rank |
| *HMG1* | S288C | 228/<br>2118 | *RIM15* | 0.12 | 2<sup>nd</sup> | 0.49 | 4<sup>th</sup> | 93 | 2<sup>nd</sup> |
|  |  |  | *CDC28* | 0.03 | 7<sup>th</sup> | 0.47 | 7<sup>th</sup> | 31 | 25<sup>th</sup> |
|  |  |  | *DBP7* | 0.11 | 3<sup>rd</sup> | 0.45 | 10<sup>th</sup> | 44 | 4<sup>th</sup> |
|  |  |  | *HMG1* | 0.03 | 9<sup>th</sup> | 0.45 | 11<sup>th</sup> | 24 | 54<sup>th</sup> |
|  | UWOPS87 | 464/<br>2556 | *RIM15* | 0.07 | 2<sup>nd</sup> | 0.42 | 15<sup>th</sup> | 66 | 2<sup>nd</sup> |
|  |  |  | *CDC28* | 0.04 | 8<sup>th</sup> | 0.44 | 4<sup>th</sup> | 53 | 3<sup>rd</sup> |
|  |  |  | *DBP7* | 0.04 | 6<sup>th</sup> | 0.40 | 44<sup>th</sup> | 36 | 14<sup>th</sup> |
|  |  |  | *HMG1* | 0.03 | 11<sup>th</sup> | 0.43 | 9<sup>th</sup> | 27 | 32<sup>nd</sup> |
|  | Y55 | 265/<br>1525 | *RIM15* | 0.09 | 2<sup>nd</sup> | 0.48 | 3<sup>rd</sup> | 55 | 2<sup>nd</sup> |
|  |  |  | *CDC28* | 0.05 | 7<sup>th</sup> | 0.46 | 8<sup>th</sup> | 37 | 7<sup>th</sup> |
|  |  |  | *DBP7* | 0.06 | 4<sup>th</sup> | 0.45 | 12<sup>th</sup> | 45 | 3<sup>rd</sup> |
|  |  |  | *HMG1* | 0.04 | 9<sup>th</sup> | 0.46 | 6<sup>th</sup> | 33 | 12<sup>th</sup> |
| *BTS1* | S288C | 224/<br>2239 | *TLG2* | 0.02 | 12<sup>th</sup> | 0.55 | 3<sup>rd</sup> | 65 | 7<sup>th</sup> |
|  | UWOPS87 | 428/<br>3743 | *TLG2* | 0.02 | 10<sup>th</sup> | 0.49 | 5<sup>th</sup> | 82 | 2<sup>nd</sup> |
|  | Y55 | 300/<br>2767 | *TLG2* | 0.04 | 5<sup>th</sup> | 0.50 | 3<sup>rd</sup> | 84 | 2<sup>nd</sup> |

### Network topology centrality analyses identify genes critical to atorvastatin sensitivity in multilayered networks

Networks may be analysed for informative topological metrics called “centralities” (Yu et al. 2007) where briefly, the more central a gene is to a network, the more biological relevance it has to the phenotype. Three centrality measurements were thus calculated separately for each GIN and PPIN as well as for the combined GIN/PPIN multi-layered network, namely betweenness centrality (shortest path length between two nodes) (Brandes 2001), closeness centrality (shortest path length between one and all other nodes) (Newman 2005), and degree centrality (number of neighbours) (Dong and Horvath 2007). As the GINs and PPINs exhibited different overall patterns of centrality (Fig EV3), the multi-layered network was prioritised to ensure consideration of both GINs and PPINs in a common analysis (Fig 8).

**Figure 8.**
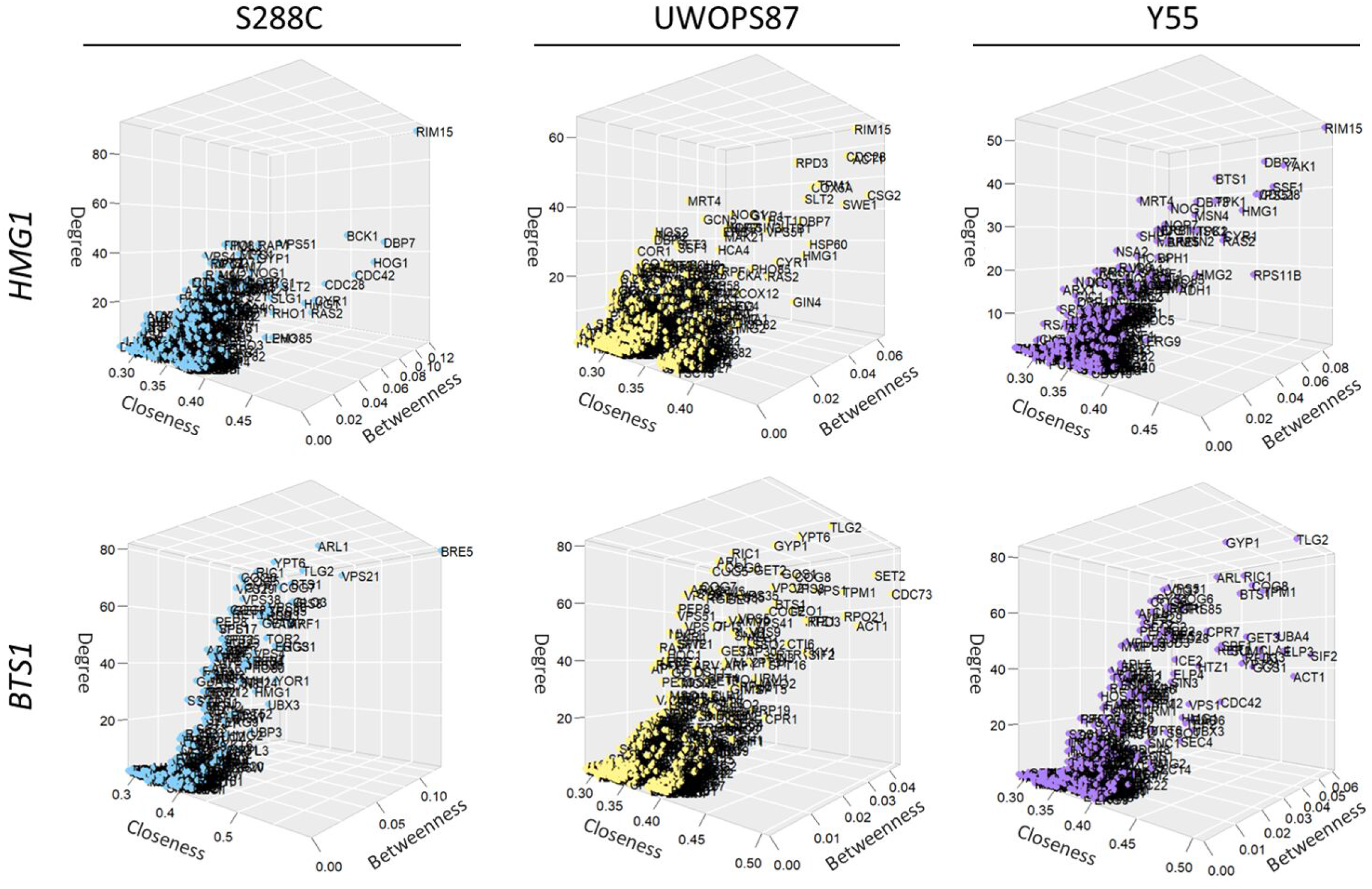
Network topology centrality analyses of multi-layered networks identify key *HMG1/BTS1* interactors for atorvastatin sensitivity. Centrality measurements (degree, closeness and betweenness) were calculated for each gene and visualised in a 3D plot. *UBI4* was excluded because owing to its highly interacting nature it skewed all the other nodes to one corner of the plot obscuring the relevance of other genes.

*RIM15*, a protein kinase involved in cell proliferation, was ranked highly in the three genetic backgrounds for betweenness, closeness and degree centralities in the *hmg1Δ xxxΔ* networks (Figs 8 and EV4; Table 1). Impressively, this result reflects the atorvastatin hypersensitivity we found for *hmg1Δ rim15Δ* (Fig 4). Betweenness, closeness and degree centrality metrics for *RIM15* were 0.12, 0.49 and 93, respectively for S288C, compared to 0.07, 0.42 and 66 for UWOPS87 and 0.09, 0.48 and 55 for Y55. Likewise, the *CDC28* kinase master regulator of mitotic and meiotic cell cycles, was also ranked highly for the three centrality metrics in the three genetic backgrounds (Fig 8; Table 1). The involvement of kinases in statin responses points to fundamental effects of statins on aspects of metabolism other than cholesterol metabolism.

For the *bts1Δ xxxΔ* networks, the t-SNARE *TLG2* that mediates the fusion of endosome-derived vesicles with the late Golgi, was distinct for the three centrality metrics in Y55 and UWOPS87 while it was less distinct in S288C (Figs 8 and EV5; Table 1). Betweenness, closeness and degree centrality metrics for *TLG2* were 0.02, 0.55 and 65, respectively for S288C, compared to 0.02, 0.49 and 82 for UWOPS87 and 0.04, 0.50 and 84 for Y55. The ubiquitin protease cofactor *BRE5* that coregulates with *UBP3* the anterograde and retrograde transport between the ER and Golgi compartments, was the highest ranked gene for the three centrality metrics in S288C, which is further supported by the atorvastatin hypersensitivity of the *bts1Δ bre5Δ* only in S288C (Fig 5). The involvement of genes mediating vesicular fusion and transport provides insight into the diverse functions of the *BTS1* branch in the mevalonate pathway.

### Community analysis identifies functional modules in multi-layered networks for three genetic backgrounds

To gain more insight into the structural organisation of the multi-layered networks in order to identify metabolic pathways mediating atorvastatin sensitivity, community analysis was conducted to partition the networks to functional subnetworks (modules) that are more interconnected compared to random (Blondel et al. 2008). We detected 3-6 modules in each network (Fig 9). Each module exhibited significant enrichment for specific metabolic pathways (*P* < 0.05), and in most cases, pathways enriched in these modules did not overlap in all three genetic backgrounds (Fig 9).

**Figure 9.**
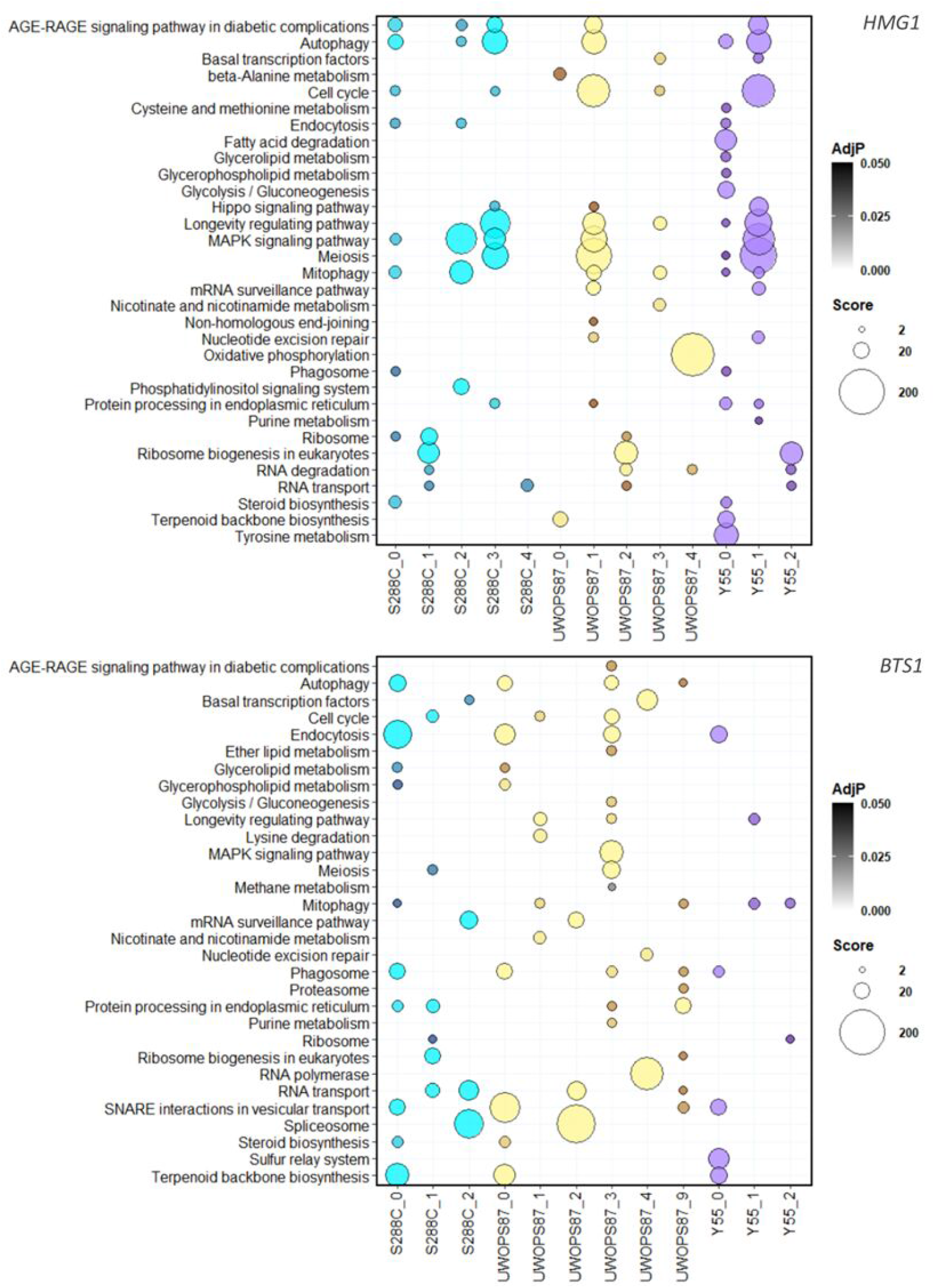
Metabolic pathway enrichment of modules in multi-layered networks for atorvastatin sensitivity. Bubble plots showing enrichment for each of the modules (named for their genetic background) identified through community analysis for *HMG1* (top panel) and *BTS1* (bottom panel) interactions. The size of the bubbles is relative to the enrichment score for each pathway, while the intensity of the colours is relative to the adjusted *P* value. The x axis labels show the genetic background followed by the number of modules. Numbers missing in the sequence are modules without significantly enriched pathways.

For the *hmg1Δ xxxΔ* networks, the longevity regulation pathway and its tightly linked processes autophagy and mitophagy were found enriched in all three genetic backgrounds (Fig 9). In contrast, the longevity regulation pathway was only enriched in UWOPS87 and Y55 for the *bts1Δ xxxΔ* networks (Fig 9). Since statins extend lifespan yet it varies person to person, and the chronological lifespan in yeast mimics the post-mitotic state of cancer cells (Bisschops et al. 2014, Jacobs et al. 2013), we sought to test the importance of specific genetic interactions to statin-mediated longevity. Consequently, we measured the chronological lifespan of *BTS1*-epistatic strains (*bts1Δ rpd3Δ, bts1Δ ras2Δ, bts1Δ rfm1Δ, bts1Δ sum1Δ, bts1Δ hst1Δ, bts1Δ sir1Δ*) representative of modules in UWOPS87 and Y55 that were enriched for the longevity regulation pathway (Fig 9). The survival integral, the area underneath the survival curve, for each single deletion strain as well as the *bts1Δ xxxΔ* strains was calculated in the presence and absence of atorvastatin over a period of 13 days (Fig 10). For S288C, the survival area remained relatively consistent across all strains with or without atorvastatin (Fig 10A), suggesting treatment did not impact the survival in this genetic background. By contrast, survival was significantly increased with treatment in UWOPS87 and was more pronounced in *bts1Δ xxxΔ* double deletion mutants compared to single deletion mutants for *sum1Δ, hst1Δ* and *sir1Δ* (Fig 10B), suggesting these specific chromatin/histone interactions with atorvastatin increase the chronological lifespan of UWOPS87 strains. For Y55, survival was significantly increased with treatment in *bts1Δ rpd3Δ* compared to *bts1Δ* and *rpd3Δ* (Fig 10C), revealing the importance of the histone deacetylase *RPD3* only in this background. These results experimentally validate the *in silico* community analyses that vary by genetic background and confirm the importance of specific chromatin/histone interactions mediating the lifespan extension activity of atorvastatin. As expected, metabolic pathways were enriched in the single-layer analysis but not in the multi-layer analysis, and vice versa (Figs EV6 and EV7), yet multi-layer networks retrieved more relevant information since GINs could not be partitioned in communities possibly due to their high connectivity.

**Figure 10.**
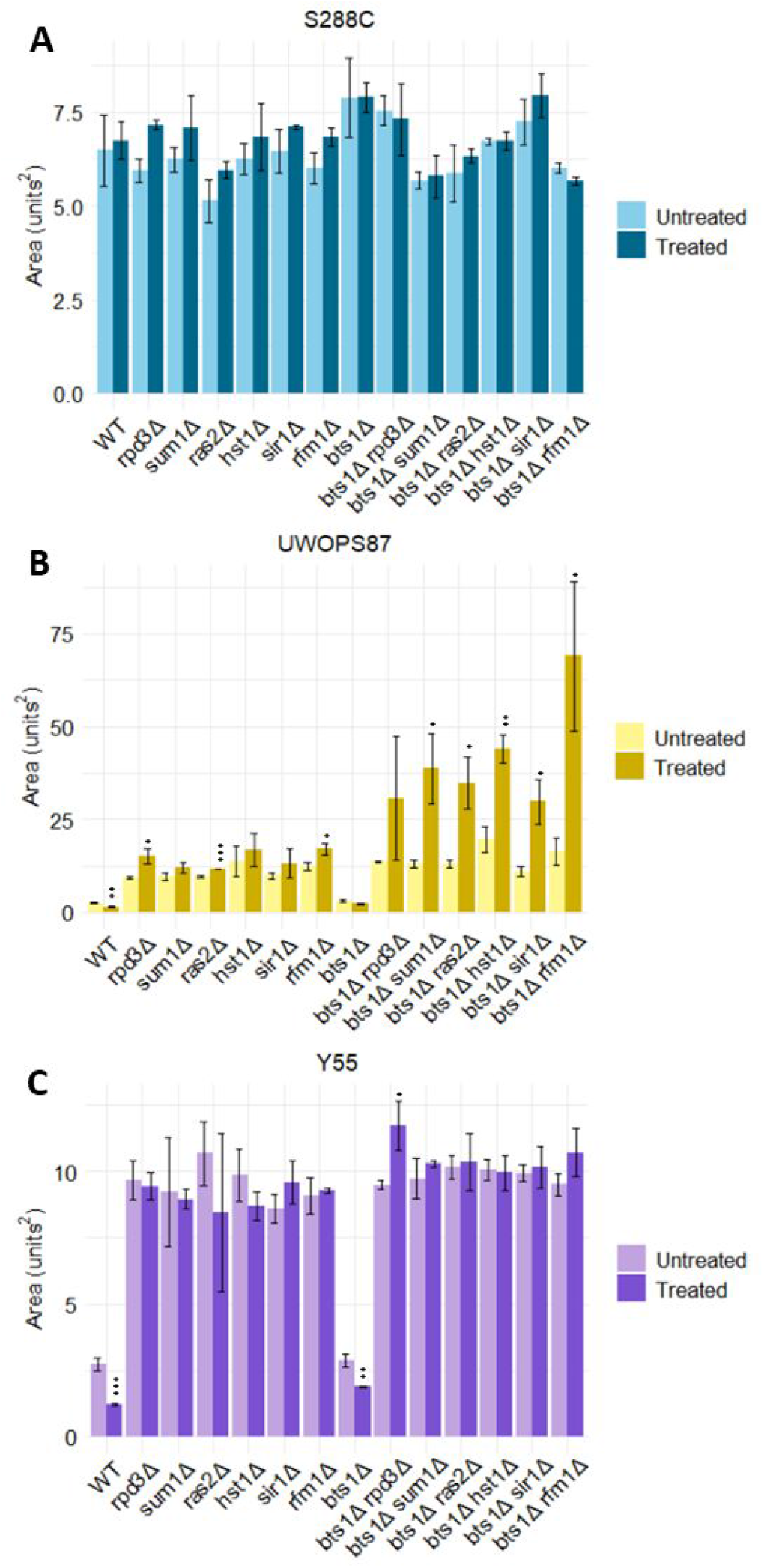
Atorvastatin treatment in the UWOPS87-2421 genetic background increases the survival integral of double mutants. Cells were grown in triplicate with and without atorvastatin. Cultures were left growing at 30°C for two weeks, and growth was measured every second day for a two-week period via hourly measurements of optical density. YODA was used to calculate surviving cell percentage. Data are shown as mean ± SD from n=3; *p ≤ 0.05, ** p ≤ 0.01, *** p ≤ 0.001, student’s t-test relative to vehicle control.

### Humanization of yeast epistasis reveals anticancer drugs for statin synergy

Combination therapies may increase efficacy of repurposed drugs (Sun et al. 2016). To see if that is the case with statins and anti-cancer drugs we identified the human orthologues of the key centrality genes identified in our yeast genomic analyses across the three genetic backgrounds (Table EV3). Since most of them have been previously annotated to cancer (Table EV1), we evaluated these genes for enrichment in the Drug Signature Database (Yoo et al. 2015) providing greater specificity in selection of synergistic combinations (Fig 11). This analysis detected 205 drugs with ‘signature genes’ integral to their bioactivity as well as atorvastatin (*P* < 0.05). Of these 205 drugs, the maximum and minimum odd ratios were 86 and 2, respectively. To compare the chemical genetic profiles of the top-ranked drugs, the odds ratio values for the top 20 drugs by *p* values and their signature genes were visualised in a bubble plot (Fig 11).

**Figure 11.**
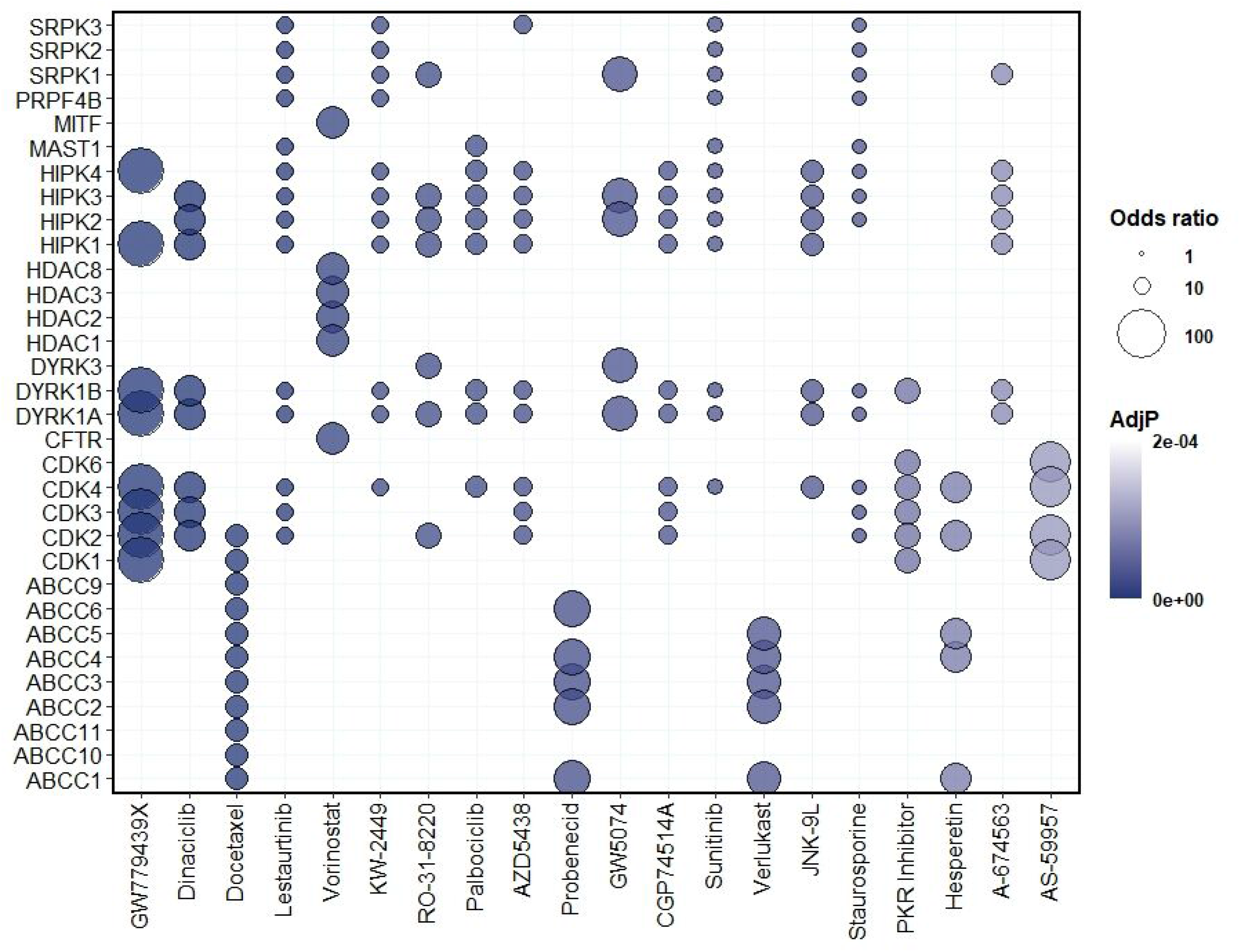
Human orthologues of yeast interactions reveal drugs/compounds to test for synergy with atorvastatin. Human orthologues of validated genes and bottleneck genes were processed via an enrichment analysis for signature genes in the Drug Signature Database. Bubble plots representing the human orthologues (y-axis) that were enriched for drugs/compounds (x-axis). The colour of each bubble is determined by the adjusted *P*-value and the size of bubble reflects a score computed by running the Fisher exact test for random gene sets to determine the deviation from the expected rank, where bigger bubbles represent greater enrichment.

The 32 signature genes shown in Fig 11 represent seven major processes targeted by specific drugs. Four drugs (docetaxel, probenecid, verlukast, hesperetin) were correlated with ABC transporter genes involved in numerous functions including drug efflux and that provoke failure of chemotherapeutics (El-Awady et al. 2016). Fifteen drugs (GW779439X, dinaclinib, docetaxel, lestaurtinib, KW-2449, RO-31-8220, palbociclib, AZD5438, CGP74514A, sunitinib, JNK-9L, staurosporine, PKR inhibitor, hesperetin and AS-59957) were correlated with kinase activity contributed by cyclin-dependent kinase (CDK) genes, dual-specificity tyrosine-regulated kinase (DYRK) genes and MAP kinase (HPK) genes involved in cell cycle. Four drugs (lestaurtinib, palbociclib, sunitinib, staurosporine) were correlated with the MAST1 gene involved in survival signalling pathways that confers cell resistance to the chemotherapeutic cisplatin (Jin et al. 2018). Four drugs (lestaurtinib, KW2449, sunitinib, staurosporine) were correlated with the PRPF4B gene, an essential gene for triple-negative breast cancer metastasis (Koedoot et al. 2019). Lastly, eight drugs (lestaurtinib, KW-2449, RO-31-8220, AZD5438, GW5074, sunitinib, staurosporine, A-674563) were correlated with serine/arginine-rich protein-specific kinase (SRPK) genes involved in activation of various signalling pathways that mediate cytotoxic effects of genotoxic agents including cisplatin (Sigala et al. 2021). The pyrazolopyridazine GW779439X ranked the highest of all drugs and compounds (*P* = 2.42E-09; odds ratio = 86), which was mainly due to key centrality genes in cyclin-dependent kinase genes (CDK genes) identified with the *HMG1* query.

Thus, the majority of the top 20 drugs (*i.e*., lowest AdjP value) identified here for potential synergy with atorvastatin have exhibited anticancer activity and only two have been investigated for statin synergy (Table EV4). These drugs with established anticancer activity include dinaciclib, docetaxel, lestaurtinib, vorinostat, palbociclib, and sunitinib. Interestingly, one of the top results is docetaxel, a well-established chemotherapeutic for the treatment of breast cancer that was previously investigated for synergy with lovastatin, albeit the trial was terminated for lack of funding (NCT00584012). Another noteworthy candidate combination therapy is probenecid, which is a drug that inhibits renal excretion and would thus increase the half-life of statin drugs. Clinical trial NCT03307252 evaluated the pharmacokinetics of probenecid with a number of drugs including rosuvastatin, but this trial did not evaluate the anticancer activity of the statin. In addition to drugs/compounds with established anticancer activity, we also propose combination therapies with GW779439X with antibiotic properties, verlukast with bronchodilator properties and hesperetin with a wide variety of properties including cholesterol-lowering, antioxidant, anti-inflammatory and anticancer properties.

## Discussion

Drug response involves many genes whose phenotypes may be the result of epistatic genetic interactions, pleiotropy and dependency on genetic background which can be analysed as multidimensional networks (Fig 2) via topological centrality metrics and community algorithms. Here, we used yeast genome-wide deletion libraries with two genetic probes (SGA query gene strains *hmg1Δ* and *bts1Δ*) and an inhibitory drug (atorvastatin) to define colony growth phenotypes and networks in the mevalonate pathway (Fig 1). Genetic background is known to affect genetic interactions (Busby et al. 2019; Deutschbauer and Davis 2005; Galardini et al. 2019), so we investigated genetic interactions in the standard S288C strain and two additional deletion libraries in the genetic backgrounds, UWOPS87 and Y55. With the network topological centrality and community algorithms used here, clear pathways of GO cellular processes emerged in the case of the *HMG1* or*BTS1* probes for interactions involved in autophagy, ageing, endocytosis, actin and unfolded protein response (UPR) pathways. The following discusses specifics of these topics.

Statins are known to activate autophagy (Mao et al. 2018; Peng et al. 2018; Okubo et al. 2020), yet the mechanism is not fully understood. Here, we distinguish *RIM15* as a key statin-modulator in positively regulating autophagy, since *RIM15*-deficiency conferred hypersensitivity in the atorvastatin-treated *HMG1* query and *RIM15* was a high betweenness gene (bottleneck) in three genetic backgrounds. Bottlenecks are of high relevance because they tend to connect functional clusters of genes (Brandes 2001; Yu et al. 2007). We enhanced the networks derived from colony growth by including published interactors with *RIM15* of pathlength 2 and 75% of the genes belonged to a single community module in all the genetic backgrounds. This community was enriched for functions involved in meiosis, longevity and autophagy. This serine/threonine kinase *RIM15* is integral to statin-induced autophagy in yeast and may be conserved in mammalian cells via the human orthologue, MASTL.

Functional redundancy (Sambamoorthy & Raman, 2018) is seen for *RIM15* in its role in actin-mediated processes and endocytosis as described here. Relatedly, we show here that *CDC28* is a top centrality gene in the *HMG1* genetic interaction networks that previously was shown to have a suppressing interaction with *RIM15* (Juanes et al. 2013; Talarek et al. 2017). *CDC28* and *RIM15* cluster together in a cochaperone module for ‘actin and morphogenesis’ (Rizzolo et al. 2017). Although *CDC28* did not belong to a statistically significant community module in our study, 86% of the genes that interacted with *CDC28* in Y55 and UWOPS87 belonged to the community module corresponding to meiosis, cell cycle and MAPK signalling, suggesting that networks are functionally redundant for these processes as well as actin/endocytosis. We note that human orthologues of *RIM15* code for cytoskeleton components, such as actin and the intermediate filament that have shown to be part of the statin response (Denoyelle et al. 2003).

*TPM1* is a bottleneck gene for UWOPS87 and Y55 treated with atorvastatin and a synthetic lethal genetic interaction with S288C in *BTS1* query. It is a major isoform of tropomyosin that binds and stabilises actin cables (Liu and Bretscher 1989). Statins have indeed shown to induce cytoskeletal reorganisation while increasing the levels of F-actin important in cancer cell motility (Sarkar et al. 2022). Pleiotropy for statins is seen where the *CDC28* human orthologue CDK1 is down-regulated by atorvastatin with anticancer activity in esophageal squamous cell carcinoma (ESCC) cells (Yuan et al. 2019). Likewise, simvastatin induced G1 arrest and inhibited cell growth of colorectal cancer cell lines by a mechanism that included down-regulating CDK4/cyclin D1 and CDK2/cyclin E1 (Chen et al. 2018). Simvastatin and lovastatin suppressed expression of CDK1, CDK2, CDK3, CDK4 and CDK6 in prostate cancer cells with reduced cell viability due to induced apoptosis and cell cycle arrest (Hoque et al. 2008). Tropomyosin is a cancer prophylaxis drug target, which could be nuanced for general toxicity by synergistic combinations of statins and drugs aimed at tropomyosin-modifying genes, prescreened for activity in the yeast models as described here. Mechanism of action provides insight into drug synergy (Cokol et al., 2011), an example here that identified probenecid for potential synergy with atorvastatin. Probenecid is prescribed for the prevention of hyperuricemia-associated gout. Given that high serum cholesterol levels have been correlated with hyperuricemia (Peng et al. 2015), it is likely that many patients worldwide are simultaneously prescribed probenecid and statins. Databases such as UK Biobank (Sudlow et al. 2015) might reveal whether simultaneous treatment of probenecid with statins has been associated with reduced rates of cancer.

Ageing is considered one of the main risk factors for cancer development (Berben et al. 2021). Here we show that double deletion of *BTS1* with *HST1* increased the chronological lifespan of the UWOPS87 strain, unlike their single deletions. *HST1*, demonstrated to be synthetic sick with *HMG1* in UWOPS87, is a NAD(+)-dependent histone deacetylase paralogue to the human SIRT1 gene sirtuin 1. Sirtuins are a family of protein deacetylases that regulate ageing and longevity (Imai and Guarente 2016; Longo and Kennedy 2006). SIRT1 has indeed been described as part of the mechanism behind the anti-ageing effect of statins (Bahrami et al. 2020; Lei et al 2014) focusing on cell senescence. However, the role of SIRT1 in the chronological ageing of non-dividing cells has not been investigated. Our results point to a genetic background-dependent role, in which the presence and potential activation of yeast *HST1* by atorvastatin does not affect chronological lifespan. However, chronological lifespan is greatly increased by its double deletion with *BTS1* in the UWOPS87 background. For the Y55 background, it was the deletion of *BTS1* with *RPD3* that increased the chronological lifespan. *RPD3* is a histone deacetylase orthologue to the human HDAC1 and HDAC2, which have been linked to the mechanism of anticancer activity of statins (Li & Gan 2017; Lin et al. 2008), and HDAC1 is known to have an anti-ageing activity in brain cells (Pao et al., 2020). We note that statins increase the lifespan of the model organism *Caenorhabditis elegans* (Jahn et al. 2020) and decreased mortality independent of cholesterol in humans aged 78-90 years old (Jacobs et al. 2013). Our results plausibly point to the importance of the *BTS1* branch with chromatin and histones in atorvastatin-mediated effects on lifespan.

Taken together, we have demonstrated the utility of using chemical genetics and multi-layered network analyses to elucidate genetic complexity of metabolic pathway phenotypes that may be behind drug molecular mechanisms. In this paper, we discuss atorvastatin and its anticancer properties. We note that UPR/ER stress is tightly linked to autophagy (Senft and Ronai 2015; Yan et al. 2015). We have previously shown UPR activation is qualitatively genetic background-dependent in response to statins (Busby et al, 2019) and ER stress is also a known mechanism of the anticancer activity of statins (Yang et al. 2010). Our model here provides more information on the link between UPR with actin-mediated endocytosis (Mattiazzi Usaj et al. 2020), autophagy and ageing (Estébanez et al. 2018; Taylor 2016). We propose that statin treatment induces the UPR, dysregulates endocytosis and causes autophagic cell death (Fig 12). It is plausible that all of these phenotypes have a role in the anticancer activity of atorvastatin via induction of UPR, especially since statins inhibit and remodel actin cytoskeleton (Boerma et al. 2008; Chubinskiy-Nadezhdin et al. 2017), actin is necessary for endocytosis (Mooren et al. 2012), and UPR is induced in yeast mutants deficient of actin-mediated steps in endocytosis (Mattiazzi Usaj et al. 2020). The many genetic interactions involving these cellular processes described here potentially provide lists for drug targets.

**Figure 12.**
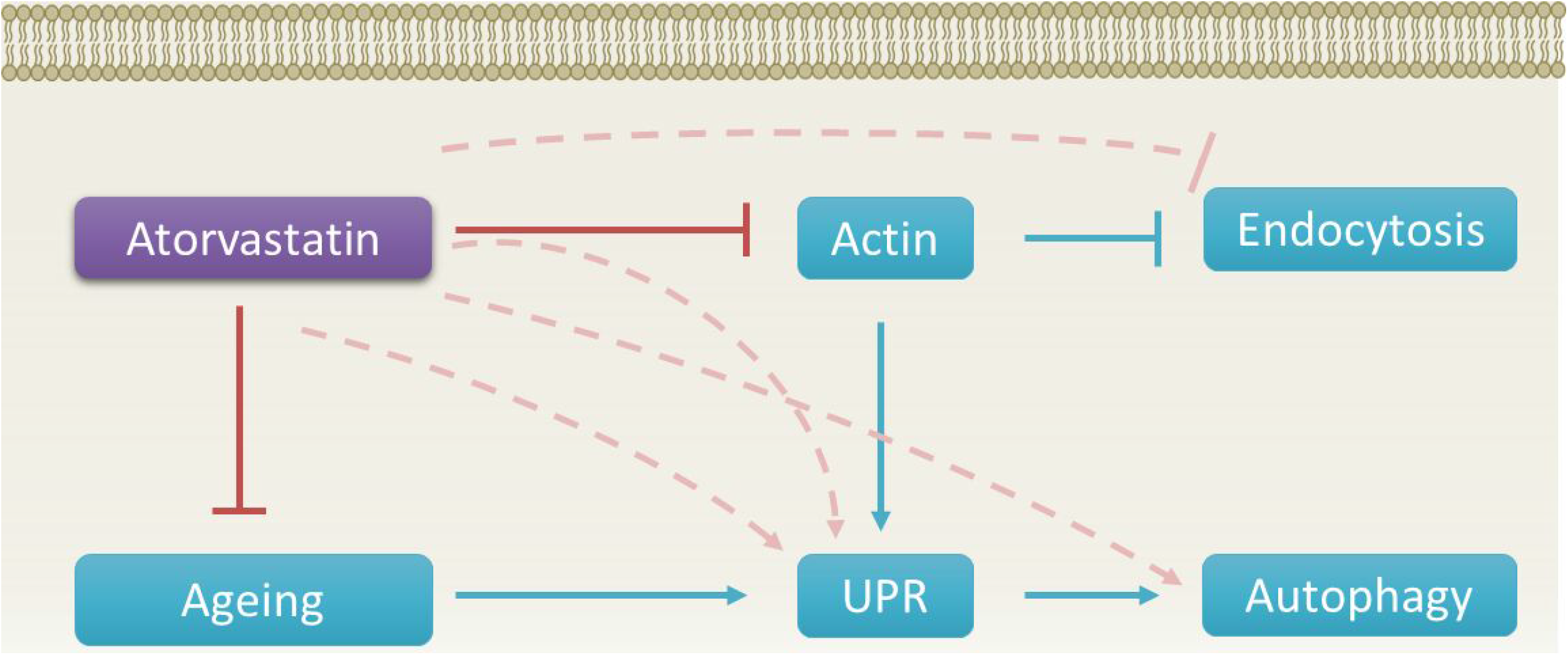
Proposed integration of mechanisms. Atorvastatin inhibits components of the actin cytoskeleton, which in turn inhibits actin-mediated endocytosis and induces UPR. Atorvastatin inhibits ageing pathways, which also results in the dual induction of UPR and autophagy. Hence, atorvastatin is an indirect inhibitor of endocytosis and indirect activator of UPR and autophagy. Red blunt head arrows point to pathways inhibited by atorvastatin. Blue arrows and blue blunt head arrow point to pathways that are inhibited or induced, respectively. Hashed pink arrows and hashed blunt head arrow point to inhibition or induction, respectively, of pathways via indirect mechanisms of atorvastatin.

## Materials and Methods

### Yeast strains, plasmids and media

The *S. cerevisiae* strains used in this study are described in Table EV5. Stocks were stored at −80°C in 15% glycerol. Strains that contained the URA3_CEN plasmid were grown on agar with 1 mg/mL of 5-Fluoroorotic Acid (5-FOA, Kaixuan Chemical Co) to select for uracil auxotrophs before construction of the query strains. DMA libraries were maintained in synthetic complete (SC), synthetic dropout (SD), enriched sporulation, or yeast peptone dextrose agar as previously described (Amberg 2005). The media and solutions used included agar, amino acids, peptone, yeast extract, yeast nitrogen base (Formedium), ampicillin, atorvastatin calcium, glucose, monosodium glutamate, potassium acetate (Sigma-Aldrich), geneticin sulfate, L-canavanine sulfate, S-aminoethyl-L-cysteine hydrochloride (thyalisine) (Carbosynth), nourseothricin sulfate (Werner BioAgents), hygromycin B (Life Technologies). All antibiotics and supplements stocks were filter sterilised with 22 *μM* pore filters (Jet Biofil).

### Synthetic Genetic Array (SGA) analysis

SGA analysis was conducted in quadruplicate as previously described (Busby et al. 2019; Tong et al. 2001), 1536-colony format in three genetic backgrounds (S288C, UWOPS87 and Y55) with newly constructed query deletion strains, in which *HMG1* and *BTS1* was replaced with NATMX antibiotic resistance gene via PCR-mediated disruption, using specific primers and cycle conditions (Table EV6). The plasmids used for this study were conserved in *Escherichia coli* (DH5α) and stored at −80°C: MX4-natR switcher cassette p4339 (Tong et al. 2001). PCR products were then transformed into a *MATα* SGA starter strain via homologous transformation as previously described (Gietz and Schiestl 2007) and integration into the genome was confirmed by PCR as previously described (Tong and Boone 2006). Plates were replica plated with an automated RoToR HDA system (Singer Instruments).

### Genome-wide growth analysis

The selected double deletion mutant libraries (*hmg1Δ xxxΔ* and *bts1Δ xxxΔ*) were pinned on SC agar, incubated at 30°C overnight, and used as an inoculum source to pin on SC agar with and without IC_30_ concentrations of atorvastatin that were determined for each genetic background. These plates were incubated at 30°C for 12 and 24 h, time points when the colonies were imaged using a digital camera (Canon). The colony sizes were quantified and scored through SGAtools (Wagih et al. 2013) where z-scores were used to compare growth with and without atorvastatin (zero indicates no difference between the control and treatment, negative scores indicate reduced fitness with atorvastatin, and positive scores indicate increased fitness with atorvastatin). All SGA scores were visualised in violin plots generated in R and based on their point of inflection, the cut-offs were selected to identify strains for experimental validation in 384-colony format and serial dilution spot assay.

### Validation of negative genetic interactions in 384-colony format

The validation of negative genetic interactions was performed in a two-step process. First, 96-colony format plates were arrayed containing no more than 29 atorvastatin-hypersensitive double mutants each with *his3*Δ control border strains and also *his3*Δ control strains surrounding each candidate to ensure the colony sizes were not biased. Each plate also included a wild type strain. The atorvastatin-hypersensitive double mutants for S288C, Y55 and UWOPS87 *hmg1Δ xxxΔ* or *bts1Δ xxxΔ* that did not overlap with the single deletions *xxxΔ* were arrayed as described. Control single deletions were also arrayed to confirm that negative interactions pertained to double deletions only. The arrayed plates were screened with the same IC_30_ concentrations of atorvastatin used in the 1536-colony format. Plates were incubated at 30°C for 24 h, imaged using a digital camera, growth was quantified using SGAtools (Wagih et al. 2013) as described above for the 1536-colony format, and hypersensitive strains were then selected for an additional experimental validation step through serial dilution spot assays.

### Validation of negative genetic interactions in serial dilution spot assay

Overnight cultures were prepared in 96-well plates and four 1:10 serial dilutions were spotted using a manual pinning tool on SC agar with and without an IC_30_ concentration of atorvastatin. Plates were incubated at 30°C for 48 h, imaged using a digital camera, and evaluated visually for atorvastatin-specific growth defects. A cut-off for growth defect was determined as one spot less of atorvastatin-treated versus non-treated strains and of the double deletion compared to the single deletions (query gene deletion and *xxxΔ*). Those atorvastatin-hypersensitive double mutants that validated in spot assays were then submitted to another round of spot assays this time including the three genetic backgrounds.

### Single-layer network analyses

Validated genetic interactions that enhanced the hypersensitivity to atorvastatin were examined in the context of gene-gene and protein-protein interaction networks. The list of validated genes was augmented with gene-gene interactions using GeneMania (Warde-Farley et al. 2010) using all available studies with a maximum number 110 interacting genes. Using NetworkAnalyst (Xia et al. 2015; Zhou et al. 2019), the list of validated genes was augmented with protein-protein interactions using the STRING database (Szklarczyk et al. 2015) that includes text-mining, genomic information, co-expression and orthology with the additional requirement for experimental evidence with a confidence score cut-off of 900. The resulting protein-protein interaction network was a first-order network representing the input nodes with their direct interactors (path length 1), which was then augmented into a second-order network to include nodes that connected the input genes as well as nodes that were interactors (of path length 2), but that only included the minimum number of nodes necessary to maintain connectivity of the network (minimum network). The gene-gene interaction networks (GINs) and the protein-protein interaction networks (PPINs) were then integrated into a single multi-layer network using TimeNexus (Pierrelée et al. 2021) in Cytoscape (Shannon et al. 2003).

### Topology centrality analysis

The single-layer and multi-layer networks were analysed for various measurements of network centrality using the NetworkAnalyzer for undirected networks application in Cytoscape (Boccaletti et al. 2014). Three centrality measurements were calculated: (1) Degree centrality, which computes the number of edges linked to each node so that a node with degree 5 has 5 edges associated, that is, it is linked to 5 other nodes (Dong and Horvath 2007); (2) closeness centrality, which corresponds to the average shortest path length of one node to every other node computed by the Newman method (Newman 2005), where 0 means an isolated node and 1 is the highest centrality and connectivity; and (3) betweenness centrality, which is the probability of passing through a node when using the shortest path length between two nodes and is computed with the highly precise algorithm developed by Brandes (Brandes 2001) to distinguish nodes critical to maintain a network. The three measurements of centrality were visualised as 3D plots using R (Soetaert 2019).

### Community analysis

Functional modules (communities) in the single-layer and multi-layer networks were determined using the InfoMap algorithm (Rosvall and Bergstrom 2008) in NetworkAnalyst (Xia et al. 2015). Statistical significance for each module was evaluated for their clustering significance or network connectivity as computed by a Wilcoxon rank-sum test (*P* < 0.05).

### Pathway enrichment analysis

Modules were investigated for their function via metabolic pathway enrichment analysis using the Kyoto Encyclopedia of Genes and Genomes (KEGG) pathway database (Kanehisa and Goto 2000) implemented in Enrichr (Chen et al. 2013; Kuleshov et al. 2016). Pathway enrichment was statistically evaluated using an adjusted *P*-value with the Benjamini-Hochberg method for correction (Benjamini and Hochberg 1995), a z-score reflecting the deviation of a Fisher’s exact test from an expected rank, and a combined score that is the product of the natural logarithm of the *P*-value multiplied by the z-score. Fold-enrichment and *P*-value (<0.05) for statistically significant pathways in each module were visualised in bubble plots using R (Wickham 2016).

### Chronological lifespan assay

The effect of atorvastatin on the chronological lifespan of *S. cerevisiae* was assessed as previously described (Murakami & Kaeberlein, 2009) with alterations. A single colony from single and double deletion strains across all three genetic backgrounds was inoculated into 5 mL of SC media and incubated overnight at 30°C with constant agitation. 50 & l of each culture was removed and added to fresh tubes with and without atorvastatin, and then incubated at 30°C with constant agitation for the course of the experiment. Outgrowth assays were conducted at various time points of 1, 3, 5, 7, 9 and 11 days where 10 & l from each culture was removed and added to a Biofill 96 well plate with 140 & l SC media. OD measurements of the plate were taken using an Envision 2102 Multilabel plate reader (Perkin Elmer) at 590 nm hourly for 48 hours while the experimental tubes were placed back in the rotator at 30°C. Data were visualised using OD as a measure of viability over time and analysed using the Yeast Outgrowth Data Analysis program (YODA) as previously described (Olsen, Murakami, & Kaeberlein, 2010) to calculate doubling time inflection, time shifts and the survival integral for each mutant and treatment over the experimental period.

### Gene set enrichment for drug signatures

Human orthologues of genes that interact with *HMG1/BTS1* query strains as well as highly ranked centrality genes were determined using Yeastmine in the Saccharomyces Genome Database (Cherry et al. 2012) and examined for significant enrichment (*P* < 0.05) in the Drug Signature Database (Yoo et al. 2015) implemented in Enrichr (Chen et al. 2013; Kuleshov et al. 2016; Xie et al. 2021).

## Acknowledgements

Funding from a Cancer Society of New Zealand Wellington Division CT Collins PhD Scholarship (to CERH) is greatly appreciated, and this research was supported in part by the Maurice Wilkins Centre for Molecular Biodiscovery Flexible Research Programme (to PHA), for which we are also very grateful. We thank Bede Busby for the Y55 and UWOPS87 deletion libraries.

## Author contributions

ABM, CERH and PHA designed the study. CERH completed all experiments and analyses except for the chronological lifespan assays completed by LJC. ABM, CERH and PHA wrote the manuscript. All authors read and approved the final version.

## Conflict of Interest

The authors declare that they have no conflict of interest.

## Data availability

The datasets produced in this study are available in Tables EV7-EV9.

## Expanded view Figure legends

Figure EV 1. **Genetic interaction networks connecting validated interactions hypersensitive to atorvastatin**. Genetic interactions networks for the *HMG1* query (upper panel) and *BTS1* query (lower panel) were constructed using GeneMania and visualised using NetworkAnalyst. Darker nodes in each network are the input genes.

Figure EV 2. **Protein-protein interaction networks connecting validated interactions hypersensitive to atorvastatin**. Protein-protein interaction networks for the *HMG1* query (upper panel) and *BTS1* query (lower panel) were constructed using STRING and visualised using NetworkAnalyst. Darker nodes in each network are the input genes.

Figure EV 3. **Network topology centrality analyses of GINs and PPINs identify key *HMG1/BTS1* interactors for atorvastatin sensitivity.** Centrality measurements (degree, closeness and betweenness) were calculated for each gene and visualised in a 3D plot.

Figure EV 4. **Metabolic pathway enrichment of modules in protein-protein interaction networks for atorvastatin sensitivity**. Bubble plots showing enrichment for each of the modules (named for their genetic background) identified through community analysis for *HMG1* (top panel) and *BTS1* (bottom panel) interactions. The size of the bubbles is relative to the enrichment score for each pathway, while the intensity of the colours is relative to the adjusted *P*-value. The x axis labels show the genetic background followed by the number of modules. Numbers missing in the sequence are modules without significantly enriched pathways.

Figure EV 5. **Network centrality of genes behind hypersensitivity to atorvastatin for *HMG1* interactors overlap in three genetic backgrounds.** Genes that ranked in the top ten centrality measurements were found to confirm phenotypic findings. Centrality measurements (betweeness, closeness and degree) were calculated in NetworkAnalyzer app in Cytoscape (Boccaletti et al. 2014) and networks were built in Cytoscape. The red outline points to highly central genes.

Figure EV 6. **Network centrality of genes behind hypersensitivity to atorvastatin for *BTS1* interactors overlap in three genetic backgrounds.** Genes that ranked in the top ten centrality measurements were found to confirm phenotypic findings. Centrality measurements (betweeness, closeness and degree) were calculated in NetworkAnalyzer app in Cytoscape (Boccaletti et al. 2014) and networks were built in Cytoscape. The red outline points to highly central genes.

Figure EV 7. **Metabolic pathway enrichment of modules that did not overlap in single-layer and multi-layer network community analysis.** Bubble plots showing enrichment for each of the modules (named for their genetic background) identified through community analysis for *HMG1* (top panel) and *BTS1* (bottom panel) interactions that were unique to either single-layer (left panel) or multi-layer (right panel) analyses. The size of the bubbles is relative to the enrichment score for each pathway, while the intensity of the colours is relative to the adjusted *P* value. The x axis labels show the genetic background followed by the number of modules.

